# Clinical Strains of *Pseudomonas aeruginosa* secret LasB protease to induce hemorrhagic diffused alveolar damage and acute lung injury in mice

**DOI:** 10.1101/2021.05.15.444319

**Authors:** Yajie Zhu, Xiaoli Ge, Di Xie, Shangyuan Wang, Feng Chen, Shuming Pan

## Abstract

**Background:** Acute lung injury and acute respiratory distress syndrome are most often caused by bacterial pneumonia, characterized by severe dyspnea and high mortality. Knowledge about the lung injury effects of current clinical bacteria strains is lacking. The aim of this study was to investigate the ability of representative pathogenic bacteria isolated from patients to cause ALI/ARDS in mice and identify the virulent factors.

**Method:** 7 major bacteria species were isolated from clinical sputum and instilled in to mice airway unilaterally. Histology study was used to judge the lung injury effect. Virulence genes were examined by PCR. Sequence type of *P. aeruginosa* strains were identified by MLST. LC-MS/MS was used to identify the suspicious protein bands. LasB was purified through DEAE-cellulose column and its toxicity was tested both in vitro and in vivo.

**Results:** *Staphylococcus aureus, Streptococcus pneumoniae, Streptococcus agalactiae, Acinetobacter baumannii, Klebsiella pneumoniae, Pseudomonas aeruginosa* and *Escherichia* coli were randomly separated and tested 3 times. Among them, gram-negative bacteria are more potential than gram-positive bacteria to cause acute lung injury. However, *P. aeruginosa* is the only pathogen which induced diffused alveolar damage, hemorrhage and hyaline membrane in the lung of mice. The lung injury effect is associated to the excreted matrix metalloproteinase LasB of *P. aeruginosa*. Purified LasB recapitulated hemorrhagic acute lung injury identical to *P. aeruginosa* infection in vivo. We found this was due to the powerful degradation effect of LasB on both lung extracellular matrix and key proteins in coagulation cascade without inducing cellular apoptosis.

**Conclusion:** *P. aeruginosa* strains are most capable to induce ALI/ARDS among major clinical pathogenic bacteria, this ability is specifically attributed to their LasB production.

## Introduction

Acute lung injury (ALI) and its more serious form acute respiratory distress syndrome (ARDS) is a major killer in the ICU. Despite decades of efforts, the pathogenesis of ALI/ARDS remains unclear, let alone effective treatment[1]. Diffused alveolar damage (DAD), hyaline membrane formation and alveolar bleeding which are common pathology features found in ALI/ARDS patients, were poorly recapitulated in animal models. Furthermore, although it is defined as a syndrome of similar clinical signs associated with damage to the alveolar–capillary membrane, the heterogenicity in ARDS have been more and more recognized [2].

The major risk factor of ARDS is pneumonia caused by various microorganisms, accounting for nearly 60% of the cases[3]. Microorganisms like bacteria or virus, evolves rapidly in response to environment for better adaption and survival [4–6]. Compared to the type strains or standard strains preserved and studied in lab for decades, clinical isolates nowadays are exhibiting significant differences in phenotype, drug resistance and invasiveness[7–9]. However, virulence of this “new generation” of bacteria in causing lung injury have not been comparatively explored.

Sputum culture are routinely prescribed for patients suspected or diagnosed with pneumonia. According to the yearly report of China Antimicrobial Surveillance Network (CHINET), *Klebsiella pneumoniae, Acinetobacter baumannii, Pseudomonas aeruginosa, Staphylococcus aureus, Streptococcus pneumoniae* and *Escherichia coli* are the 6 species most commonly found, accounting for 70.5% of the 90955 strains separated from clinical respiratory specimens nationwide in 2020. Coinfection of different bacteria species is usual in pneumonia patients, sometimes even secondary to virus or fungus infection[10, 11]. Are these pathogenic bacteria equally virulent in causing lung tissue injury? This led us to wonder perhaps there is one or some bacteria particularly have dominated in the pathogenesis of ALI or even ARDS.

As Xinhua hospital is a member of CHINET[12], the bacteria prevalence here is a representative part of the status in China. In this study, we selected to isolate the 6 major bacteria species listed above plus *Streptococcus agalactiae* from clinical sputum, based on both the data from CHINET and our own hospital. Since the most relevant features of ALI/ARDS in animal models are rapid onset (within 24 h) and histological evidence of tissue injury[13], our research mainly applied histopathology methods to visualize lung injury 24 h post treatment. To better illustrate the effect of each bacteria, we applied a unilateral lung injury model by inoculating microorganisms specifically into the left lung while keeping right lung unharmed as control. This relieved dyspnea, allowed mice to survive until the intended time and formed a natural comparison within the individual for comparison.

We found that among all the species, *P. aeruginosa* is the only bacteria caused classic ALI/ARDS pathology change in mice, characterized by severe hemorrhage, diffused alveolar damage and hyaline membrane formation. We discovered this is caused by the LasB elastase (pseudolysin) which degraded the lung matrix and key proteins in coagulation pathway.

## Materials and methods

### Ethical statement

All experiments referring to the use of animals in this study were approved by the Institutional Animal Care and Use Committee of Shanghai Xinhua Hospital affiliated to Shanghai Jiao Tong University School of Medicine (XHEC-F-2018-047).

### Animals

Pathogen-free C57BL/6 mice (female, 10 weeks old, 21±1g), SD rats (female, 6-8 weeks old, 200±10g) were purchased from Shanghai Sippr-BK laboratory animal Co. Ltd. All animals were accommodated at the Model Animal Research Center of Xinhua Hospital in a specific pathogen-free animal facility under constant temperature and humidity, with sufficient qualified food and water for 1 weeks before use.

### Unilateral intratracheal instillation

In mouse, we select the left lung for bacteria or bacteria exoproducts administration. Meanwhile, the right lung was set as the control lung within each individual. To perform unilateral lung intubation, a 24G, 19mm long intravenous indwelling catheter (Introcan, BRAUN) was used as the intratracheal tube and inserted through a cervical incision. Mice were anesthetized by 1% pentobarbital sodium (50mg/Kg) intraperitoneally and put to a proper supine position. The anterior cervical hair was removed and the skin disinfected with 75% ethanol. In order to expose the trachea, a small longitudinal incision was made. The indwelling catheter was then punctured gently into the trachea between the second and third cricoid cartilage. As anatomical data shows that the mouse main trachea is approximately 10mm long, therefore the 19mm cannula full length inserted will enter the left or right bronchus. Left bronchus intubation was achieved when the insertion angle is 5-10° to the anterior median line as indicated in Figure 1A. Then, 25 μl bacteria suspension or 30 μl P. aeruginosa exoproducts were deposited into the left lung of each mouse using microliter syringes. After instillation, mice were placed on a warm blanket, resting to left recumbent position until fully awake.

**Figure 1.**
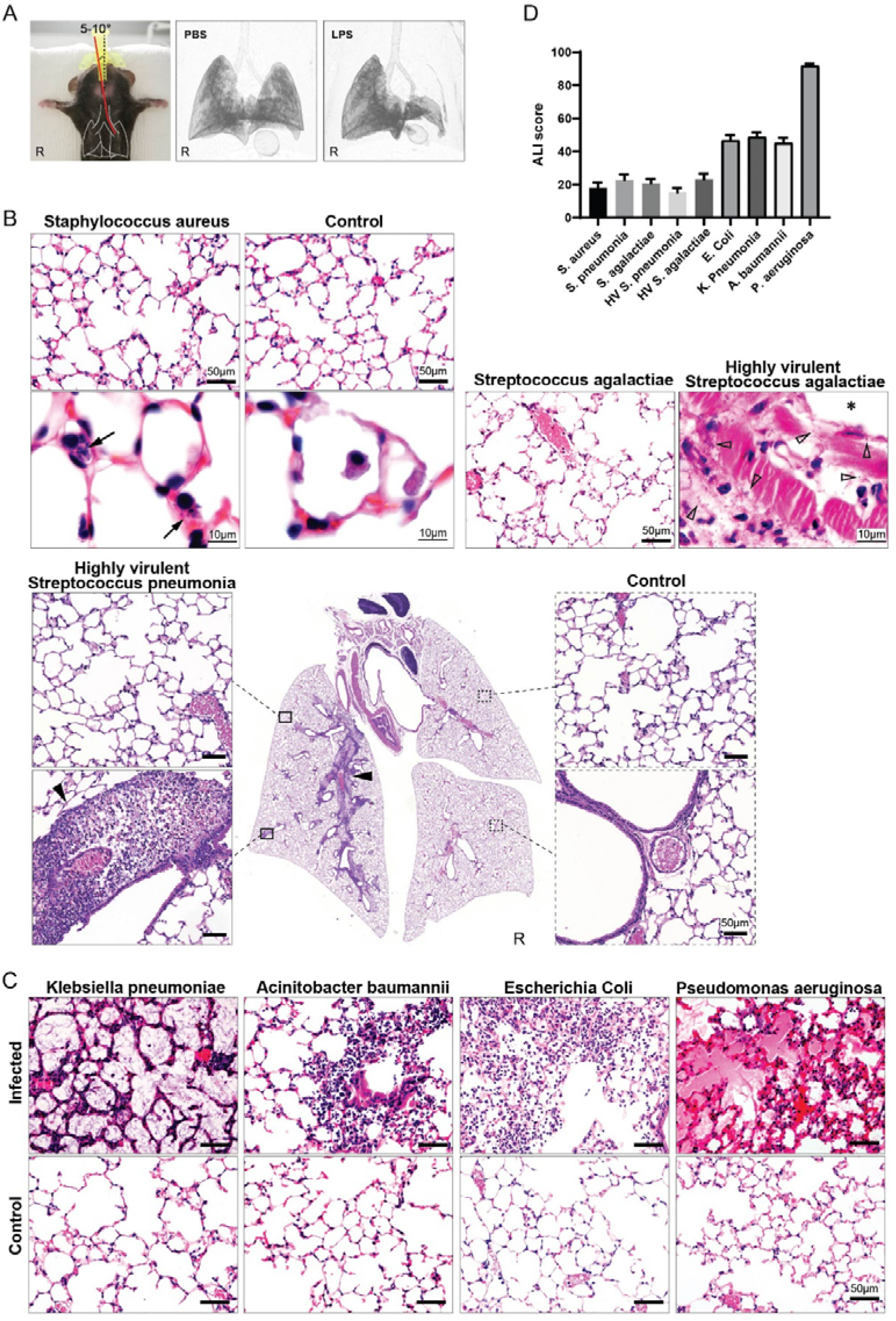
Histological assessment of acute lung injury induced by representative bacteria separated from clinical sputum. (A) Endotracheal intubation. Unilateral lung infiltration 4 hours after LPS instillation (3D CT reconstruction). (B) Hematoxylin and eosin (HE) stain of lung sections of gram-positive bacterial infection. *S. aureus*, *S. agalactiae* or *S. pneumonia* suspension was instilled unilaterally. Clean alveolar, no neutrophil infiltration noted in 24 h. Coccus found in lung macrophages (black arrows). In high virulent gram-positive strains lung infection, large amount of coccus in the perivascular space (black arrowheads) or migrating into the vascular lumen (*) were seen (empty arrowheads), without alveolar damage. (C) HE stain of lung sections inoculated with gram-negative bacteria. Alveolar injury occurred in all G-bacteria infected lungs. *K. pneumoniae*: neutrophils in the alveolar wall and K. pneumoniae filled the alveolar space; *A. baumannii*: patchy neutrophil infiltration; *E. coli*: diffused neutrophil infiltration; *P. aeruginosa*: hemorrhage, hyaline membrane, vessel congestion, alveolar wall thickening and neutrophil infiltration. (D) Score of different bacteria induced ALI 24h post inoculation.

### Lung histopathology

Mice were sacrificed at 24h post-infection. The lung was inflated with 4% paraformaldehyde fixative under constant pressure of 15 to 25 cm H2O via the trachea by standard procedures[14]. Lung and heart were harvested en bloc and submerged in fixative for approximately 24 hours before the heart was removed. The lung was embedded in paraffin block for tissue sections (5 μm). Hematoxylin and eosin staining or TUNEL staining were performed subsequently. ALI score was performed as described[13]. The modified ALI score substituted the item “Neutrophils in the interstitial space” to “Red blood cells in the alveolar space” to better differentiate the severity of lung injury resulted from P. aeruginosa exoproducts.

### Clinical isolates

Major bacteria species were separated from clinical sputum specimen based on morphology, confirmed by mass spectrometry and stored at -70℃. For enrichment culture, non-fastidious bacteria: *Staphylococcus aureus, Streptococcus agalactiae, Acinetobacter baumannii, Klebsiella pneumoniae, Pseudomonas aeruginosa* and *Escherichia coli* were inoculated into LB broth (Beyotime, China) incubated overnight at 36±1°C, 60rpm. Fastidious bacteria: *Streptococcus pneumoniae* was cultured on sheep blood agar plates (Yi Hua, Shanghai) overnight at 36±1°C, 5%CO2. Bacteria from exponential growth were collected, washed twice and resuspended in sterile normal saline (NS). The turbidity was adjusted to 3 McFarland standards (MCF), which equals 9×108 CFU/ml, by the standard procedure for later use. Each of the 7 species was isolated and prepared as above three times randomly.

### *P. aeruginosa* strains and exoproducts

A total of 8 *P. aeruginosa* strains were randomly isolated from clinical respiratory tract specimen. They are named as Pa 1-6 according to the separation time, plus a mucoid strain named as Pa M and a strain used in the earlier experiments as Pa J (isolated in July). The growth medium for routine harvesting of *P. aeruginosa* exoproducts was Columbia agar supplied with 5% sheep blood (90 mm, Comagal, Shanghai). The entire agar surface was inoculated with a cotton swab. Plates were incubated at 36±1°C for 3 days. Bacteria and exoproducts on the plate were dissolved in sterile NS (5 ml per plate). Bacteria cells were removed after centrifugal for 3200 × g, 30 min and further cleared by 0.22 μm filters. Then the exoproducts acquired was concentrated by ultra-centrifugal filters at 3000 × g, 30 min (10 kDa cutoff value, Amicon Ultra, Merck). Protein level was measured through BCA method. All exoproducts were adjusted to the same protein concentration by adding sterile NS.

### PCR and MLST

Virulence genes were detected by PCR. Briefly, a single colony of *P. aeruginosa* was added to 1ml ddH2O, washed twice and resuspended. We extracted bacterial DNA by boiling the suspension for 10 min at 100 °C. Using pairs of specific primers (Table S1), we examined the presence or absence of the following genes among clinical isolates of *P. aeruginosa*: aprA, toxA, lasA, lasB, plcH, prpL, exoS, exoY, exoT, exoU, lepA, phzS, rhlR, mucA, algR and fliC (Table 1). PCR amplifications were performed in 50 μl reaction mixtures. Based on product size (Table S1), genes were identified and recorded as positive (+) or negative (-).

**Table 1.**
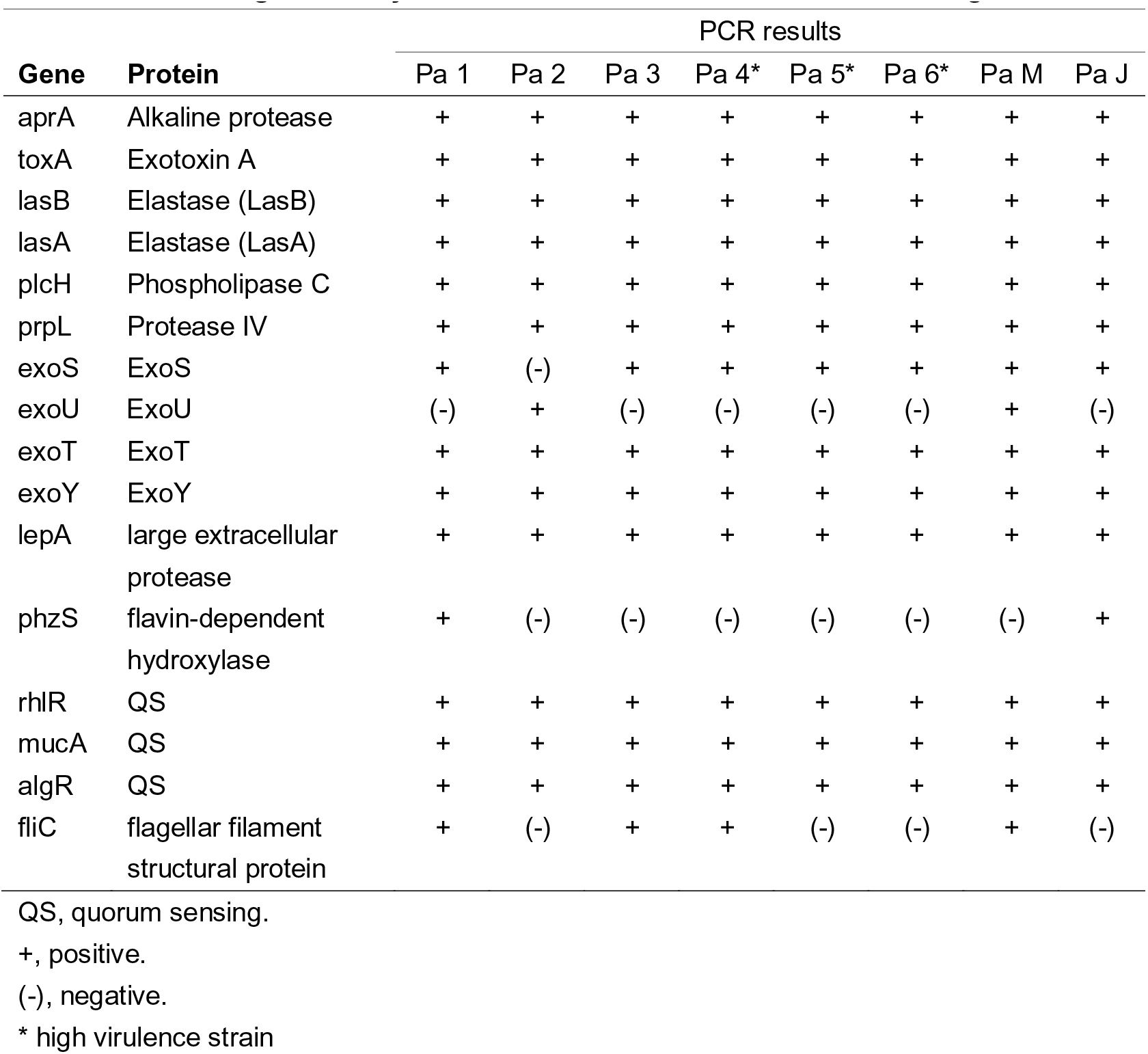
Virulence genes analysis in clinical isolates of Pseudomonas aeruginosa.

For multilocus sequence typing, seven house-keeping genes: acsA, aroE, guaA, mutL, nuoD, ppsA, trpE were amplified and sequenced according to standard procedures[15](Sangon, Shanghai). Allele type and sequence type were acquired by querying on PubMLST website[16].

### SDS-PAGE

10% gel was used for SDS-PAGE of exoproducts or proteins under 150V constant voltage.

### Gelatin zymography

Gelatin zymography was performed as reported by Kessler and colleges[17]. 8% gel containing 0.1% gelatin was used in SDS-PAGE. P. aeruginosa exoproducts (2mg protein/ml) were preincubated with 25 mM EDTA or NS for 30 min at room temperature. 3.5 μg of each sample (measured by protein) was mixed with non-reducing loading buffer (1% SDS added), without boiling, loaded at interval wells to avoid cross-contamination. Electrophorese was performed at 4 °C, 110 V constant voltage. The gels were washed twice before transferred to incubate within 200ml development buffer (Tris-HCl 50 mM pH 7.5, CaCl2 5mM, NaCl 0.2 M, Brij-35 0.02%) for 16–20 h at 37 °C. At last, gels were stained by Coomassie blue dye R-250 (0.25%) for 1 h, decolorized by destainer (methanol 20%, acetic acid 10%, 70%ddH_2_O) until clear bands were seen, and scanned by Amersham Imager (AI680, GE).

### Protease assays

*P. aeruginosa* exoproducts and purified LasB were tested for elastase activity using Elastin-Congo red (Sigma) method[17]. EnzChek Elastase Assay Kit (Thermo Fisher Scientific) was used to monitor the proteolytic activity of eluted fractions in chromatography process according to the manufacturer’s instructions.

### Fibrinogenolytic and fibrinolytic activities

Fibrinogenolytic activity was assayed using bovine fibrinogen as a substrate. Exoproducts of Pa 1-6, M, J (1 mg protein/ml) and porcine elastase (50 U/ml) were preincubated with protease inhibitor cocktail (17 mM AEBSF, 2.5 μM Aprotinin, 1 mM Bestatin, 0.1 mM E64 and 0.1 mM Leupeptin in DMSO) or EDTA (8.5 mM) at room temperature for 30 min. 100 μl of fibrinogen solution (4 mg/ml in saline) was mixed with 10 μl of each sample and incubated at 37 °C for 1 h. 5 μl of the product from each sample was subjected to SDS-PAGE.

For the fibrinolytic activity, 50 μl fibrinogen (6 mg/ml in PBS) and 30 units of thrombin were mixed in a 200 μl PCR tube, and preincubated for 15 min at 37 °C to form the fibrin clot. Then, 15 μl exoproducts (1 mg protein/ml) were added to each tube and incubated for 1 h at 37 °C. The sample was gently stirred using a syringe needle to check for fibrin fibers, and the solubility of fibrin was observed.

### Thrombin degradation activity

Exoproducts of Pa 1-6, M and J (1 mg protein/ml) were preincubated with 100 mM EDTA at room temperature for 30 min. 2 of each sample was mixed with 100 μl bovine thrombin solution (2 mg/ml) and incubated at 37 °C for 1 h. 5μl of the mixture was subjected to SDS-PAGE. The thrombin activity was also measured, by mix 5μl of each product with 50 μl fibrinogen (6 mg/ml in PBS), incubated at 37 °C for 15 min and then checked for fibrin fibers formation.

### Proteomic analysis

Exoproducts from different strains were separated on 10% polyacrylamide gel electrophoresis (SDS-PAGE) and stained with Coomassie blue. The whole lane or certain protein bands of interest were excised from the gel and processed by mass spectrometer (LTQ Orbitrap Velos Pro, Thermo Finnigan) coupled to liquid chromatography as described[18]. Mascot 2.3 software was used for data analysis.

### Purification of LasB

The protease was purified by ammonium sulphate precipitation followed by DEAE-cellulose chromatography. *P. aeruginosa* strain Pa 4 was cultivated on Columbia agar plates (90mm) supplied with 5% sheep blood at 37℃ for 3 days (For each chromatography column we use 8 plates). Each plate was washed by 8-10ml 20 mM Tris-Cl buffer (pH 8.0). After centrifugated for 3200 × g, 60 minutes, the pellet (bacterial cells) was discarded. Ammonium sulphate was added to the supernatant up to 20% saturation, sediments removed by centrifugation at 3200 × g for 30 min, and continued up to 80% saturation, precipitates were collected by the same way and resuspended in 20 mM Tris-Cl buffer (pH 7.5). This crude enzyme solution was desalted using ultra centrifugal filters of 10 kDa cutoff value (Amicon Ultra, Merck) and then loaded on a 15 mm× 200 mm DEAE cellulose column pre-equilibrated with 20 mM Tris-Cl buffer (pH 7.5). After washing the column with the same buffer, elastase was eluted by applying a linear gradient of NaCl from 0 to 1000 mM at a flow rate of 0.34 ml/min. Fractions of 2.5 ml were collected. Activity of the fractions were measured using EnzCheck Elastase Assay Kit (Thermo Fisher). Active fractions were concentrated by ultracentrifugation described as above and stored -20℃ for later use.

### Cytotoxicity assay

THP-1 cells were seeded in 96-well plates with 1640 medium containing 10% fetal bovine serum. Purified LasB in concentration of 100 μg/ml, Pa 4 exoproducts in concentration of 200 μg/ml or Pa 4 bacteria 7.5×10^7^ CFU/ml were added to wells. Cytotoxicity was measured by LDH release 4 hours post incubation using a LDH cytotoxicity assay kits (Beyotime Biotechnology) according to the manufacturer’s instruction.

### Hemostasis function

To examine the effect on hemostasis function, 20 μg LasB was added to 2ml blood (1.8ml blood + 0.2ml sodium citrate anticoagulant) from healthy volunteers, incubated for 35 min at room temperature. The samples were then subjected to automatic hemostasis tests by ACL TOP (550 CTS, Werfen).

### Statistics

The data were expressed as the mean ± standard error of mean. Statistical analysis was performed in the GraphPad Prism 8 software using one-way or two-way t-test or log rank test as appropriate. Statistical significance is indicated as (*, P < 0.05; **, P < 0.005; ***, P < 0.0005). Data are representative of at least three independent experiments.

## Results

### Unilateral lung injury in mice

We used a unilateral lung injury model in this study through tracheotomy and intubation into the left lung of the mice. We verified the method through LPS instillation. As shown by CT scan (3D reconstruction) in Figure 1A, the instillation of 30 μl 5mg/ml LPS successfully induced consolidation of the whole left lung in 4 hours.

This animal model was initially designed to see if heavy unilateral lung injury would provoke bilateral alveolar damage through systemic impacts like cytokine storm, whereas, we could find no such phenomenon in mice tested with LPS, acid, porcine elastase or mechanical ventilation, even if severe inflammation and animal death were observed (unpublished data). The untreated lung in unilaterally injured mice exhibited no difference to the lung from intact normal mice (Figure S1A).

### Gram-positive bacteria lung infection

2.25×10^7^ CFU of *S. aureus*, *S. agalactiae* or *S. pneumoniae* suspended in 25μl PBS were instilled into the left lung of mice to study the lung injury effect in vivo. Much to our surprise, all mice groups remained healthy 24 h post inoculation, and had no symptoms of acute lung injury such as dyspnea, weight loss or reduced activity. Similar in all three gram-positive bacteria infected groups, lung histology found no visible alveolar damage. We did find a few coccus in the in the infected lung in contrast with the control lung, however, they were located in the cytoplasm of alveolar macrophages (Figure 1B), suggesting the infection were already in control of the immune cells.

Considering the daily isolated gram-positive bacteria may have relatively low virulence, we further tested a highly virulent mucoid serotype 3 *S. pneumoniae* strain and a hypervirulent *S. agalactiae* strain isolated earlier, of which we had to reduce the bacteria number to 2.25×10^5^ CFU and 1.5×10^7^ CFU respectively to keep the inoculated mice live to 24 h endpoint. However, no visible alveolar damage occurred either in both groups after treatment, even though 50% mice died within 24 hours or developed seizure, sepsis and multiple organ infection (Figure S1B). Neutrophil accumulation, red blood cell effusion and large amount of coccus appeared in the interstitial space of pulmonary arteries, however, without provoking inflammation in the alveolar space (Figure 1B).

These results imply that irrelevant to their virulence, the gram-positive bacteria isolated from clinical sputum evoked a lung injury pattern more similar to bronchial pneumonia rather than the diffused alveolar damage found in ALI/ARDS.

### Gram-negative bacteria lung infection

Next, we tested the pathogenicity of the major gram-negative bacteria species, which together account for 60% of all clinical respiratory isolates. Mice were instilled with 25 μl bacteria suspensions (2.25×10^7^ CFU) of *Klebsiella pneumoniae*, *Acinetobacter baumannii*, *Escherichia coli* or *Pseudomonas aeruginosa*. This time, alveolar injury of different degrees occurred in all animals 24 hours after inoculation.

Interestingly, the lung injury caused by *K. pneumoniae* was noted for large number of leukocytes accumulated in the alveolar wall while the alveolar space filled with mucus, which was later proved to be all *K. pneumoniae* with thick capsular structure (Figure 1 C &Figure S1C). We don’t know yet if infection of *K. pneumoniae* had disabled the infiltration ability of neutrophils or the bacteria had digested the neutrophils infiltrated. Empyema was found in 80% of mice (n=10), probably due to the fact that *K. pneumoniae* isolated in our hospital for recent years are mainly hypermucoviscous, i.e., hypervirulent type.

Lungs infected with *A. baumannii* and *E. coli* both showed evidence of alveolar inflammation. *E. coli* triggered more diffused neutrophil accumulation in the alveolar space than the patchy pattern in *A. baumannii*. However, no septal thickening, hyaline membrane formation or red blood cell effusion were found in both bacteria.

Strikingly, *P. aeruginosa* strains from 3 independent isolation all caused hemorrhage, severe hyaline membrane formation, neutrophilic infiltration, parenchymal edema and proteinaceous debris deposit in the infected lungs, fitting with the histologic change of diffused alveolar damage.

We then compared the ALI score of each species. Generally, gram-negative bacteria achieved higher score than gram-positive bacteria, indicating they are stronger pulmonary damaging pathogens. Distinguished from all bacteria species, *P. aeruginosa* is the one and only one acquired the highest score, suggesting its great potence in ALI/ARDS pathogenesis. To know if this injury effect is a common feature or by chance, we further isolated more *P. aeruginosa* strains.

### ALI by strains of *P. aeruginosa* in vivo

To examine the virulence in vivo, 8 *P. aeruginosa* strains: Pa 1-6, Pa M and Pa J were stilled in mouse airways and survival curves were established. Quality control strain ATCC27853 was used as the control. Surprisingly, all *P. aeruginosa* strains induced hemorrhagic diffused alveolar damage in vivo (Figure 2C, Figure S2A). ATCC27853 is the least virulent in all, and Pa 3 is the least virulent among clinical isolates (Figure 2A). Pa 2 is less toxic than Pa 4, 5, 6 (p=0.003), but more toxic than Pa 1, 3, M and J (p < 0.0001).

**Figure 2.**
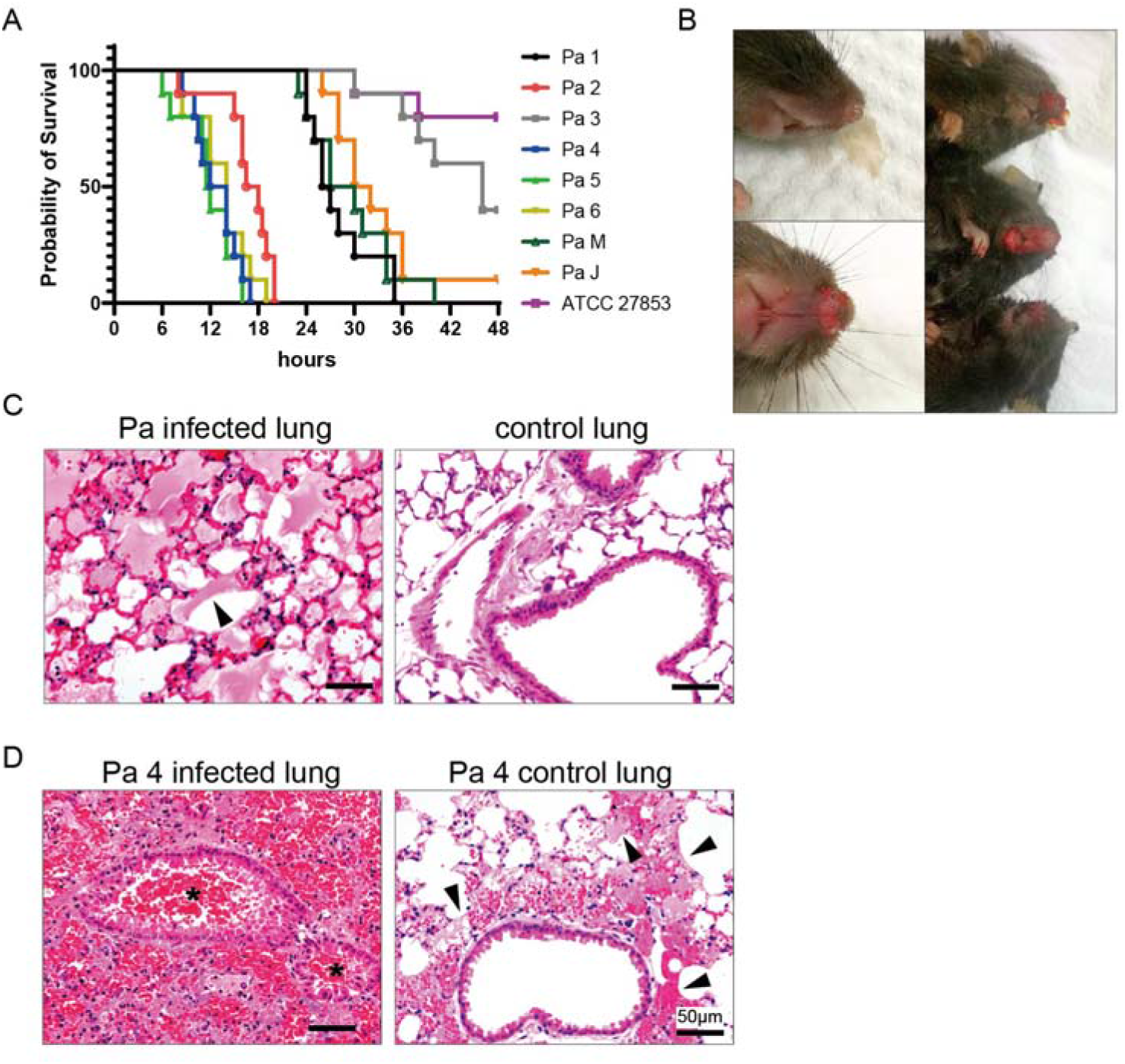
ALI/ARDS induced by different strains of *P. aeruginosa* in mice. (A) Kaplan-Meyer survival curves for mice infected with different strains (n=10 in each group). (B) Pink frothy sputum and bloody sputum from the nose and mouth of mice observed in Pa 2, 4, 5 and 6 infected group. (C) Representative histology finding in average *P. aeruginosa* lung infection: hemorrhage, neutrophil infiltration, hyaline membrane formation (arrowheads) and septal wall thickening in the alveolar. HE stain; (D) Representative histology finding in Pa 4, 5, 6 infected lungs: severe hemorrhage, alveolar structure destruction and bilateral infiltration. Bronchia lumen (*) inundated with red blood cells. Multiple hyaline membrane formation (arrowheads), diffused alveolar bleeding and neutrophil accumulation in less harmed control lung. HE stain;

Mice instilled with Pa 2 and Pa 4, 5, 6 all died within 24 h. Pink frothy sputum or bloody sputum were seen excreting from the nose and mouth in 50% of the animals (Figure 2B). Autopsy found strain Pa 4, 5, 6 caused similar pathology change prominent with lung hemorrhage. Alveolar were inundated by red blood cells, accompanied with structural destruction, neutrophils infiltration and septal thickening (Figure 2D). Also, 87% mice infected unilaterally with these 3 strains developed bilateral lung injury, suggesting that Pa 4, 5, 6 are highly deleterious. However, in the uninoculated lungs, hyaline membrane formation and red blood cell effusion were mostly observed around the bronchus or bronchia instead of alveoli, indicating the injury was likely been caused by aspiration of the bloody sputum or bacteria from the infected lung.

### Virulence genes

*P. aeruginosa* are known for owning a plethora of virulence factors, represented by T1SS, T2SS, T3SS, T6SS and quorum sensing system. T1SS and T2SS exert wide range damaging effect through secreted toxin A or proteases. Conversely, T3SS and T6SS inject several toxins into nearby cells directly. To explore the molecular mechanisms of P. aeruginosa pathogenicity, we set up PCR tests to examine the possible virulence genes. All strains are positive with toxA, aprA, lasB, lasA, plcH, prpL, lepA which are genes most related to T1SS and T2SS. Differences were detected in T3SS (exoS and exoT) and bacterial structure genes, however, the differences are not related to their level of virulence tested earlier (Table 1).

We then compared the antibiotic resistance and pyocyanin production (Table 2). Pa 2 showed resistance to Cefoperazone-sulbactam, Meropenem and Levofloxacin, while fortunately, Pa 4, 5, 6 were still sensitive to all three antibiotics, meaning their virulence were not related to drug resistant ability either. We also ruled out pyocyanin, for only Pa 1 and Pa J produced pyocyanin.

**Table 2.** Drug resistance and other characters of P. aeruginosa isolates.

|  | Pa 1 | Pa 2 | Pa 3 | Pa 4 | Pa 5 | Pa 6 | Pa M | Pa J |
| --- | --- | --- | --- | --- | --- | --- | --- | --- |
| **Antimicrobial agents** |  |  |  |  |  |  |  |  |
| Cefoperazone-sul<br>bactam | S | R | S | S | S | S | S | I |
| Meropenem | S | R | S | S | S | S | S | S |
| Levofloxacin | R | R | S | S | S | S | S | I |
| **Pigment producing** |  |  |  |  |  |  |  |  |
| Pyocyanin | + | (-) | (-) | (-) | (-) | (-) | (-) | + |

S, susceptible; I, intermediate; R, resistance.
+, positive.
(-), negative or absent.

Judged from morphology (Figure S2B), Pa 4, 5, and 6 were very similar. To know if they were actually the same, we performed multilocus sequence typing (MLST) on 8 isolates. However, all 8 isolates have different sequence types. From them, four new sequence type of P. aeruginosa: strain Pa 3, 5, 6 and M were identified. A new allele type of ppsA from Pa 3 was found (Table 3). Allelic profiles and the ST types of the isolates are shown in Table 3.

**Table 3.** Multilocus sequence typing of clinical isolates of *P. aeruginosa*.

|  | ST | Allele types |  |  |  |  |  |  |
| --- | --- | --- | --- | --- | --- | --- | --- | --- |
|  |  | acsA | aroE | guaA | mutL | nuoD | ppsA | trpE |
| Pa 1 | 3405 | 17 | 5 | 5 | 4 | 137 | 4 | 3 |
| Pa 2 | 1076 | 5 | 4 | 57 | 62 | 1 | 1 | 26 |
| Pa 3 | 3580* | 15 | 5 | 36 | 11 | 27 | 189* | 2 |
| Pa 4 | 3118 | 6 | 5 | 11 | 11 | 4 | 4 | 193 |
| Pa 5 | 3576* | 5 | 4 | 57 | 62 | 4 | 1 | 26 |
| Pa 6 | 3577* | 1 | 178 | 26 | 3 | 1 | 4 | 34 |
| Pa M | 3578* | 6 | 5 | 11 | 4 | 4 | 3 | 3 |
| Pa J | 871 | 16 | 3 | 1 | 5 | 1 | 55 | 61 |

ST, sequence type.
\* newly found in this research.

### Exoproducts induced ALI

Enlightened by the results of PCR, we decided to test the virulence of the secreted exoproducts from *P. aeruginosa*. As shown in Figure 3A, the exoproducts reproduced the typical histologic change of *P. aeruginosa* infection in lung. After instilled with high concentration of exoproducts collected from 8 *P. aeruginosa* strains (4 mg protein/ml), severe hemorrhage and DAD were observed in all mice (n=3 mice, 8 group). To our surprise, 14 of 24 mice died within 3 hours. Autopsy found tension pneumothorax and atelectasis of the whole left lung in them. On the contrary, survived mice presented emphysematous alveolar destruction, suggesting pneumothorax caused by exoproducts was most likely responsible for the sudden death in mice.

**Figure 3.**
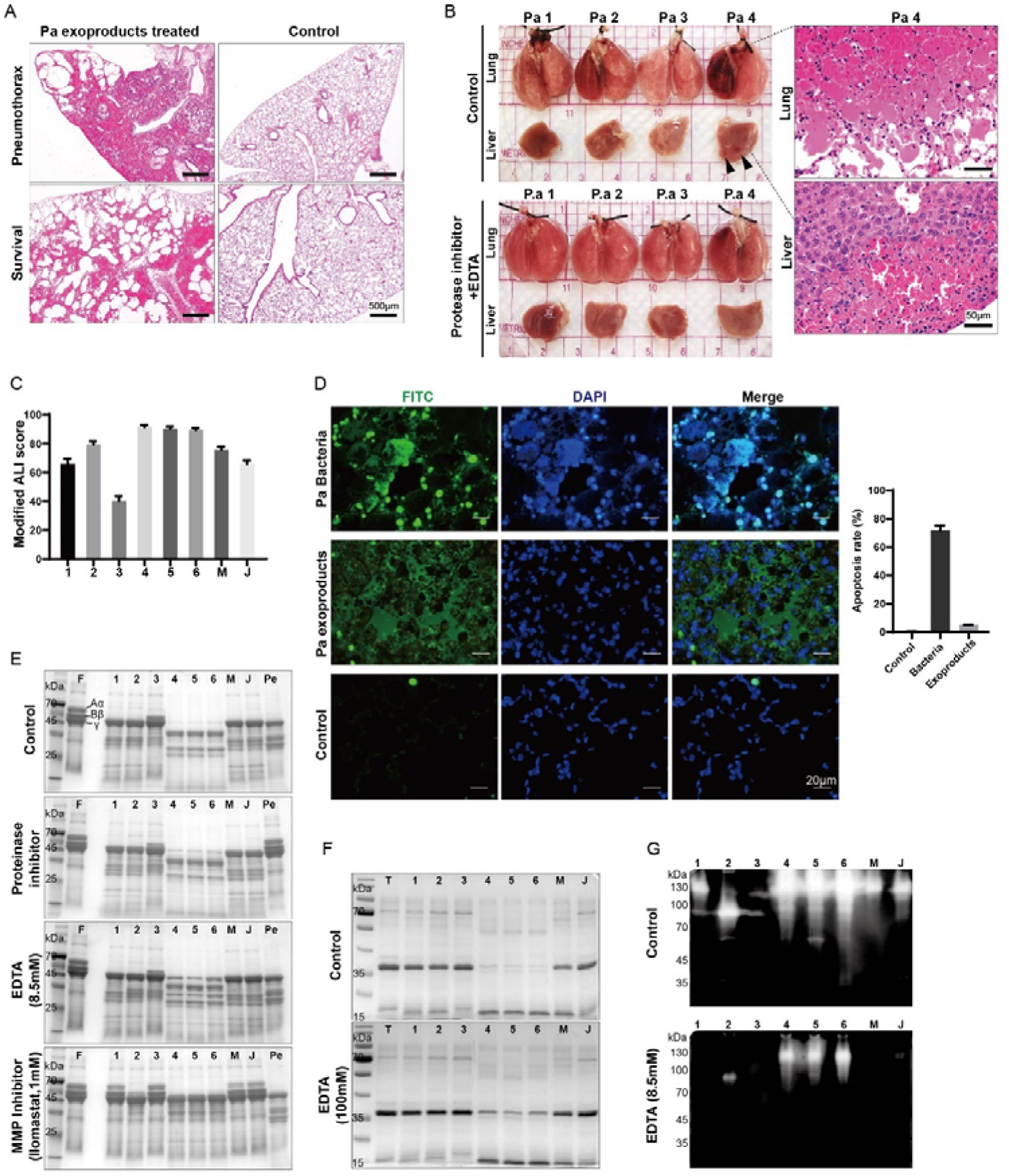
*P. aeruginosa* exoproducts induced hemorrhagic ALI in mice. (A) Diversified histology presentation caused by high concentration of exoproducts instillation. Bullae formation, severe bleeding and consolidation of the treated lung from mice died of tension pneumothorax, compared with emphysematous alveolar destruction and hemorrhage in survived mice. (B) Representative gross picture of *P. aeruginosa* exoproducts injured mice lung. Unilateral lung injury largely inhibited by protease inhibitor cocktail with EDTA added. Severe hyaline membrane, diffused alveolar damage, lung hemorrhage and subcapsular liver hemorrhage formed (arrowheads) in Pa 4 exoproducts intratracheal instilled mice. (C) ALI score of different *P. aeruginosa* strain exoproducts challenged lungs. (D) Representative TUNEL stain of *P. aeruginosa* bacteria/exoproducts challenged lungs. (E) Fibrinogen degradation effect of exoproducts. F, fibrinogen; Pe, porcine elastase (F) Thrombin degradation effect of exoproducts. T: thrombin. (G) Gelatin zymography of exoproducts.

Interestingly, after been heated at 100℃ for 10 minutes, the exoproducts lost the ability to cause alveolar hemorrhage or destruction, but only evoked neutrophil infiltration (Figure S3A). Similarly, the bacterial composition extracted from *P. aeruginosa* strains induced neutrophil accumulation but no hemorrhage (Figure S3B). Based on these evidences, we speculate the virulence factors are most likely proteins, especially exotoxin A or proteinases.

To avoid pneumothorax, we reduced the exoproducts to 1/4 (1 mg protein/ml) of the concentration used in Figure 3A. Mice were sacrificed after 24 h. Still, heavy bleeding and hyaline membrane occurred in all animals. The severity was different, judged from gross appearance of left lungs (Figure 3B). Exoproducts of Pa 4 instilled lung had also induced subcapsular liver hemorrhage. A combination of proteinase inhibitor cocktail with EDTA added largely alleviated the injury. ALI score shows clearly that exoproducts from Pa 4, 5, 6 are the strongest in induce lung injury, while Pa 3 is the weakest (Figure 3C). This tendency is in comply with the virulence of the corresponding bacteria (Figure 3D).

To find which pulmonary cell is the main victim, TUNEL labeling were used to stain both *P. aeruginosa* bacteria and exoproducts damaged lungs (Figure 3D). To our surprise, although both caused severe DAD, the exoproducts only induced 5.19% pulmonary cells apoptosis compared to the 71.7% cellular apoptosis rate caused by live bacteria. This means cell damage is not essential for the appearance of DAD.

Consequently, we infer the hemorrhagic lung injury may have resulted from the disturbance of hemostasis and destruction of extracellular matrix. We then examined the proteolysis ability of exoproducts to the two key proteins in clotting process: fibrinogen and thrombin. Figure 3E shows the fibrinogenolytic activity of the exoproducts compared with porcine elastase. The Aα and Bβ chains of fibrinogen were hydrolyzed by all exoproducts in 1 h. However, Pa 4-6 exoproducts totally degraded the γ chain of fibrinogen as well. The proteolysis was suppressed by EDTA, and to a great extent by matrix metalloprotease inhibitor (Ilomastat), but was not suppressed by proteinase inhibitors, suggesting the virulence effectors are most likely matrix metalloproteases. The freshly formed fibrin clots were also digested by exoproducts (data not shown).

Thrombin degradation assay exhibited a similar result (Figure 3F). However, the EDTA usage was much higher due to the Ca^2+^ ions contained in the commercially acquired thrombin.

Collagen and elastin are the major protein composition of pulmonary matrix. They provide mechanical stability and elastic recoil, which are essential for physiological lung function and initiation of biochemical signals in extrinsic blood coagulation. To visualize the degradation of collagen, gelatin zymography method were used (Figure 3G). All exoproducts from *P. aeruginosa* exhibited bright bands on zymography, especially Pa 4, 5 and 6. Distinguished from the rest, Pa 2 exoproducts showed a major bright band near 80 kDa, whereas the other exoproducts manifested mainly at the same level at 120 kDa. The activity can be mostly inhibited by EDTA.

We then measured the elastin lytic activity of each exoproducts using elastin-Congo red method (Table 4). All exoproducts have elastinolytic activity, which level has a tendency to be positively related to the ALI score of exoproducts and the virulence of the corresponding *P. aeruginosa* strains. The results reveal the exoproducts from *P. aeruginosa* are powerful in destroying pulmonary structural compositions, especially Pa 4, 5 and 6.

**Table 4.** Elastase activity of purified LasB and exoproducts measured by Elastin-Congo assay.

| Purified LasB | Crude enzyme | Pa 1 | Pa 2 | Pa 3 | Pa 4 | Pa 5 | Pa 6 | Pa M | Pa J |
| --- | --- | --- | --- | --- | --- | --- | --- | --- | --- |
| 90.96 | 4.225 | 1.59 | 3.23 | 0.57 | 12.5 | 12.6 | 15.1 | 3.33 | 2.18 |

### Identification and purification of virulent protein from exoproducts

Exoproducts from 8 *P. aeruginosa* strains were subjected to SDS-PAGE to analyze the secretory proteome. The gel was stained by Coomassie blue (Figure 4A). Protein bands of interest were excised carefully and subjected to LC-MS/MS analysis. The intensive band appeared at the molecular mass of 70 kDa from less virulent strains first caught our attention, which however were found to be catalase or acetylated catalase. Then we identified the most abundant protein secreted by hypervirulent strains Pa 4, 5 and 6 (at 35 kDa) as LasB elastase. Further analysis showed relative LasB expression in exoproducts of *P. aeruginosa* strains resembles their lung damage ability (Figure 4B).

**Figure 4.**
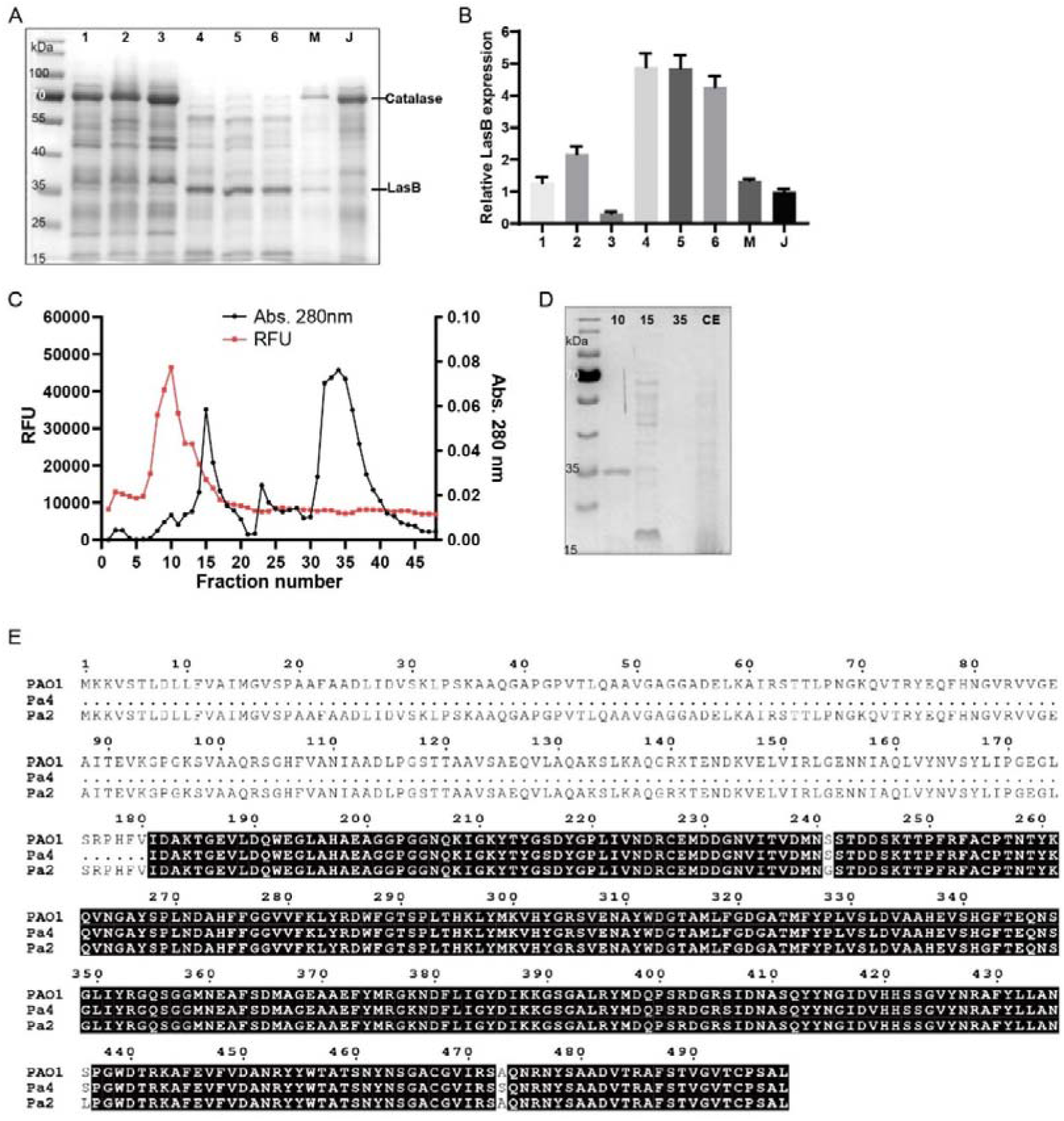
LasB elastase identification and purification. (A) SDS-PAGE of exoproducts from 8 *P. aeruginosa* clinical isolates. (B) Relative LasB expression of 8 clinical isolates. (C) Elution profile of LasB protease through DEAE 52 cellulose column. RFU: relative fluorescence unit. (D)Verification of LasB purity by SDS-PAGE. 10, purified LasB (Fraction 10); 15, impurities (Fraction 15); 35, impurities (Fraction 35); CE, crude enzyme solution. (E) Amino acid sequence alignment of Las B from Pa 4, Pa 2 and PAO1.

As it was shown in gelatin zymography, Pa 2 presented a major band at 80 kDa, different to others. However, LC-MS/MS analysis of the band found no other proteases than LasB (Figure 4E). Meanwhile, no LasA was detected from Pa 2 or Pa 4.

Taken together the results of LC-MS/MS and gelatin zymography, we selected strain Pa 4 to culture and harvest LasB. LasB was separated from the exoproducts by DEAE-cellulose. As shown in the elution profile (Figure 4C), the only active enzyme was eluted between 0.1 M to 0.25 M NaCl (fractions 5-15). Fraction 10 was collected and the protein contained was concentrated to 2mg/ml. The purity was verified by SDS-PAGE as a single band at 35 kDa, and confirmed to be LasB elastase (Figure 4D). Purified LasB exhibited an increase in elastase specific activity measured by elastin-Congo red assay (Table 4).

### LasB elastase induced hemorrhagic acute lung injury through degradation of alveolar matrix and key proteins in coagulation cascade

To confirm LasB is the effector in *P. aeruginosa* exoproducts provoked acute lung injury, 30 μl of LasB ranging from 0.005 to 2 μg/μl were tested in the left lungs of mice. Significant DAD, hemorrhage, hyaline membrane and emphysematous alveolar destruction were clearly formed by LasB ranging from 0.6 μg to 6 μg (Figure 5A). The injury could be suppressed by EDTA, MMP and EDTA plus low concentration of protease inhibitor cocktail (Figure 5B). However, the damage was aggravated by high concentration of protease inhibitor cocktail, which further tests found the cocktail itself, when instilled in high concentration, could cause rapid death in mice without alveolar tissue damage. We don’t know why yet. Meanwhile, we found the LasB dosage higher than 6 μg caused tension pneumothorax and death in 70% of the mice (n=10).

**Figure 5.**
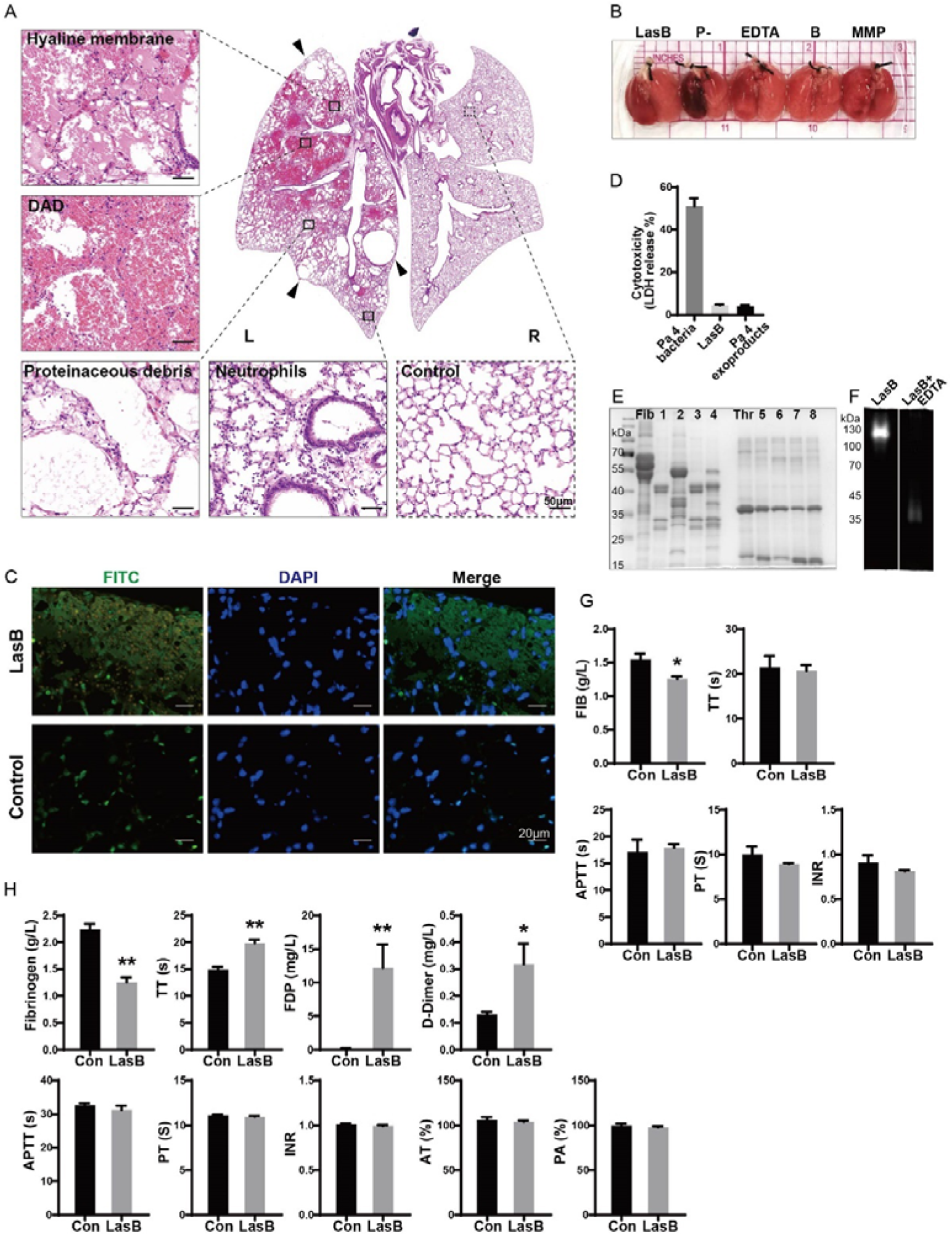
LasB elastase induced hemorrhagic acute lung injury through degradation of alveolar matrix and key proteins in coagulation cascade. (A) HE stain of mice lung unilaterally instilled with 3 μg LasB elastase. Apparent destruction to the whole left lung. Interwoven injury pattern of DAD, hemorrhage, hyaline membrane, alveolar wall thickening, proteinaceous debris, neutrophils accumulation and multiple bullae formation (arrowheads); L, left; R, right. (B) Gross picture of lungs injured by LasB and effects of different inhibitors. LasB: LasB elastase, P-: protease inhibitor cocktail; EDTA: EDTA (25mM); B: 1/2 protease inhibitor cocktail + EDTA (12.5mM); MMP: Ilomastat (1mM). (C) TUNEL stain of LasB injured lungs. No apparent lung parenchymal cells apoptosis. (D) Cytotoxicity assay of LasB or Pa 4 bacteria to THP-1 cells. (E) Degradation of fibrinogen and thrombin with or without inhibitors in vitro. Fib, fibrinogen; 1: LasB + fibrinogen; 2: LasB + Ilomastat + fibrinogen; 3: LasB + TLCK + fibrinogen; 4: LasB + TLCK + fibrinogen; Thr, thrombin; 5: LasB + thrombin; 6: LasB + Ilomastat + thrombin; 7: LasB + TLCK + thrombin; 8: LasB + TLCK + thrombin; (F) The impact of LasB on coagulation function. LasB elastase and blood from healthy donors co-incubated for 35 min, room temperature. (G) Clotting function in rats 1 h after injection of LasB elastase from tail vein.

Same as the exoproducts, TUNEL stain detected no significant difference in cellular apoptosis between LasB damaged lungs and intact lungs (Figure 5C), meaning LasB has little cytotoxicity to pulmonary cells in vivo. Also, THP-1 cytotoxicity assay found purified LasB was not virulent to cells, unlike the bacteria (Figure 5C).

However, LasB exhibited strong ability to degrade key enzymes in clotting process. After incubated at 37 ℃ for 1 hour, 12.5 μg/ml LasB hydrolyzed the 4mg/ml fibrinogen totally and the 2mg/ml thrombin partially (Figure 5E). EDTA and Ilnomastat both suppressed the process. On the contrary, TLCK exerted no inhibitory effect. This helped to rule out the trace of Protease IV.

Purified LasB appeared as a single clear band at molecular weight of 120 kDa on gelatin zymography, proved its ability to cleavage collagen. Interestingly, after been incubated with EDTA, it only left a vague band near 35 kDa (Figure 5F).

From the fibrinogen degradation assay, we have noticed that purified LasB as well as Pa 4, 5, 6 exoproducts caused a thoroughly fibrinogen degradation. We speculate this might lead to an increase in plasma fibrinogen degradation products (FDP) and D-Dimer. We thus added LasB to human blood drawn from health donors. Results are shown in Figure 5H. LasB had drastically increased FDP and D-Dimer level and elongated thrombin time (TT), causing DIC like coagulopathy in vitro. In addition, it didn’t change the indicators of endogenous coagulation pathway such as APTT, PT INR, nor affected the activity of antithrombin (AT) or plasminogen (PA). Also, we studied on if it degrades fibrinogen in vivo by injecting LasB intravenously to rats (2mg/kg). Significant decline of fibrinogen was observed within 1 h (Figure 5G). However, no difference detected in thrombin time in vivo.

## Discussion

Pneumonia is the major risk factor of ALI/ARDS. In most patients suspected or diagnosed of pneumonia, several bacteria could be found coexisting in the sputum. In this study, we examined the ability of these pathogenic bacteria isolated from patients to induce ALI in mice. Among 7 major species, gram-positive bacteria didn’t cause obvious alveolar injury, even the hypervirulent ones. In comparison, gram-negative bacteria are much more deleterious in lung. *P. aeruginosa* was found to be the most virulent, inducing hemorrhagic diffused alveolar damage, pink frothy sputum and death in animals within a few hours. We found the ALI score is positively related to their LasB elastase level. Purified LasB per se sufficiently induced diffused alveolar damage and tissue bleeding in mice. High dosage of LasB could even cause spontaneous tension pneumothorax, bullae formation and death of mice in merely 3 hours. However, our results show LasB exhibited little cytotoxicity to pulmonary cells. Further exploration validated this LasB metalloprotease is extremely efficient to hydrolysis the main component of lung structure and key proteins in clotting process such as collagen, elastin, fibrinogen, fibrin and thrombin. Collectively, our results suggest that by inducing matrix degradation and coagulopathy, LasB is the most important virulence effector of clinical *P. aeruginosa* isolates in lung infection.

*P. aeruginosa* is an opportunistic pathogen ubiquitously presenting in the environment. Despite the significant changes of microbial spectrum of infections in intensive care, *P. aeruginosa* has held a nearly unchanged position in the rank order of pathogens causing ICU-related infections for more than 4 decades[19]. Multiple resistance mechanisms enabled this microorganism to escape a wide range of antibiotics. Multidrug-resistant (MDR) *P. aeruginosa* has aroused serious concerns over its close relation to high mortality of patients with hospital-acquired and ventilator-associated pneumonia (VAP) in the ICUs[20].

Among the arsenal of virulence factors armed by *P. aeruginosa*, type III secretion system (T3SS) is characterized as the major one responsible for alveolar epithelial injury in patients. T3SS could directly inject four toxins: ExoU, ExoT, ExoS and ExoY into the cytosol of target eukaryotic cells when in contact. Strains of *P. aeruginosa* possessing the exoU gene are considered most deleterious [21]. However, among the 8 strains we collected, exoU+ genotype Pa 2 is not as virulent as exoU-genotype Pa 4, Pa 5 and Pa 6 in causing hemorrhagic ALI, indicating that T3SS is probably not as important in ALI/ARDS pathogenesis.

Exoproducts of *P. aeruginosa* are composed of many cell-associated and extracellular virulence factors, such as LPS, flagellin, proteases, exotoxins, pyocyanin, siderophores, hemolysins, and phospholipases[22]. Among them, we found ALI is mainly caused by proteases. However, *P. aeruginosa* secrets at least six proteases including alkaline protease (AprA), elastase A (LasA), elastase B (LasB), large exoprotease A (LepA), protease IV (PIV) and Pseudomonas small protease (PASP). Many researches had found *P. aeruginosa* proteases caused severe lung hemorrhage and injury in animals, however, the effect was not specifically attributed to the LasB elastase, as opposed to LasA elastase or other proteases [23–27]. Moreover, most studies investigated the role of LasB in lung infection used defined deletion or insertion mutants of the LasB gene, or used broad-spectrum metalloprotease inhibitors alternatively, which may not be assigned unequivocally to the impact of LasB [28–30]. Instead, our team successfully purified LasB elastase from a highly lung destructive *P. aeruginosa* strain and elucidated the direct causative effect of LasB in both vitro and vivo.

Further more, we ruled out the contribution of LasA, LepA, PASP and alkaline protease in ALI due to the fact the LC-MS/MS detected no LasA elastase, LepA or PASP from strain Pa 4 exoproducts, while both SDS-PAGE and zymography of purified LasB showed no bands at the molecular weight fits to alkaline protease as reported[17]. We applied TLCK inhibition assay to further ruled out the participation of PIV. Our study strongly suggest LasB is the most important lung injury protease secreted from *P. aeruginosa*.

*P. aeruginosa* LasB elastase (pseudolysin) is a neutral metalloprotease with both proteolytic and elastolytic activities that could be inhibited by metal chelators such as EDTA or MMP inhibitors. LasB has been shown to degrade a vast array of host proteins including structural components such as elastin/ collagen/laminin, immune factors like IgA/IgG/TNF-α/ IFN-γ/IL-2/IL-6/α1-antiprotease and proteins important in coagulation function like fibrin/fibrinogen/thrombin[28]. Likewise, we have validated that the important protein components in lung and clotting process can be efficiently hydrolyzed by LasB.

We also tested the commercially available recombinant LasB protein produced by *E. coli* (30R-3401, Fitzgerald) in pilot studies, however, it showed no elastolysis in vitro nor lung destruction in vivo, indicating that the toxicity of LasB mainly attributes to its activity than immunogenicity (data not shown).

Even though LasB is sufficient to provoke all necessary histopathological change identical to DAD found in ALI/ARDS, it didn’t show obvious cytotoxicity to THP-1 cells or to pulmonary cells. Jose at al. had reported similar results, that moderate LasB protease is nontoxic to Hep2 cells[31]. However, in real *P. aeruginosa* infection, not only DAD, but also extended cell apoptosis were observed. Since both LasB and exoproducts have shown no obvious cytotoxicity, we suppose the apoptosis might be caused by the other virulent factors such as T3SS and T6SS, which are highly efficient, but requires direct bacteria-to-cell contact[21]. Also, we found the most abundant protein secreted by less virulent *P. aeruginosa* strains is catalase, which probably explains their clinical prevalence, for catalase offers strong protection over nutrient-starved and biofilm bacteria from H_2_O_2_ and antibiotic-mediated killing[32].

The exoproducts of Pa2 manifested a major band near 80 kDa compared with other exoproducts (near 120 kDa) on gelatin zymography, however, the LC-MS/MS analysis of the band showed no other proteases than the neutral metalloproteinase LasB. This is in compliance with the report from Marquart et al.[33]. We infer the Pa 2 produced LasB might have some difference in polymer conformation since the aggregation of LasB monomers to different polymer forms on zymograms is commonly found[34].

FDPs and D-Dimers are the products of degraded fibrinogen or fibrin. As LasB exhibited strong ability to decompose fibrinogen, we wondered if it would raise the level of FDP and D-Dimer. Incubation of LasB with heathy donors’ blood confirmed our hypothesis, for only 35 mins, it significantly decreased fibrinogen level and drastically increased both FDP and D-Dimer level, causing DIC-like clotting disfunction. Interestingly, LasB didn’t disrupt anticoagulant process. This may aggravate the coagulopathy even more, and helped to explain the heavy bleeding in lung. LasB administrated intravenously in rat has decreased fibrinogen level in vivo as well. Whereas, thrombin time elongation was not found. We are not sure if this was caused by the strong compensatory capacity of the SD rats or by some other reasons yet. More exploration will focus on this matter in the future.

In this study, we isolated 8 clinical strains of *P. aeruginosa*, they are different in many aspects such as morphology, pigment production or T3SS genes. LasB production is the one of the few common features they share and correlates in proportion with their virulence in lung. Not only *P. aeruginosa*, a large variety of microorganisms secret extracellular proteinases too, like collagenase from *Clostridium histolyticum* and thermolysin from *Bacillus thermoproteolyticus* [35]. *S. aureus, S. agalactiae* and *S. pneumoniae* are known for their hyaluronidase production [36]. These enzymes enabled bacteria with extraordinary power to break barriers and spread [35, 37]. However, due limited time and resource, we haven’t been able to test and identify the virulence factor of every clinical isolate. Also, as our research focused on the major respiratory tract isolates and some subtypes in China, it is not known whether our results are fully applicable to other areas, where bacterial prevalence pattern may be quite different. Further studies are needed.

The research of Zupetic. J and colleges have showed that *P. aeruginosa* elastase activity was common in ICU respiratory isolates representing 75% of samples and was associated with increased 30-day mortality[38]. In this study, we demonstrated that *P. aeruginosa* is the queen of pulmonary pathogens. Among all its exoproducts, LasB per se is sufficient and necessary to elicit hemorrhagic diffused alveolar damage in animals. Monitoring LasB levels might be helpful in predicting the outcome of ALI/ARDS patients.

## Acknowledgements

We would like to thank Dr. Jingxian Liu (Division of Medical Microbiology, Department of Clinical Laboratory, Xinhua Hospital) as well as Professor Yong Zhang (Department of Immunology, SJTUSM) and Professor Min Wang (Department of Histology and Embryology, SJTUSM) for their skilled advice and assistance;

---

**Figure S1.**
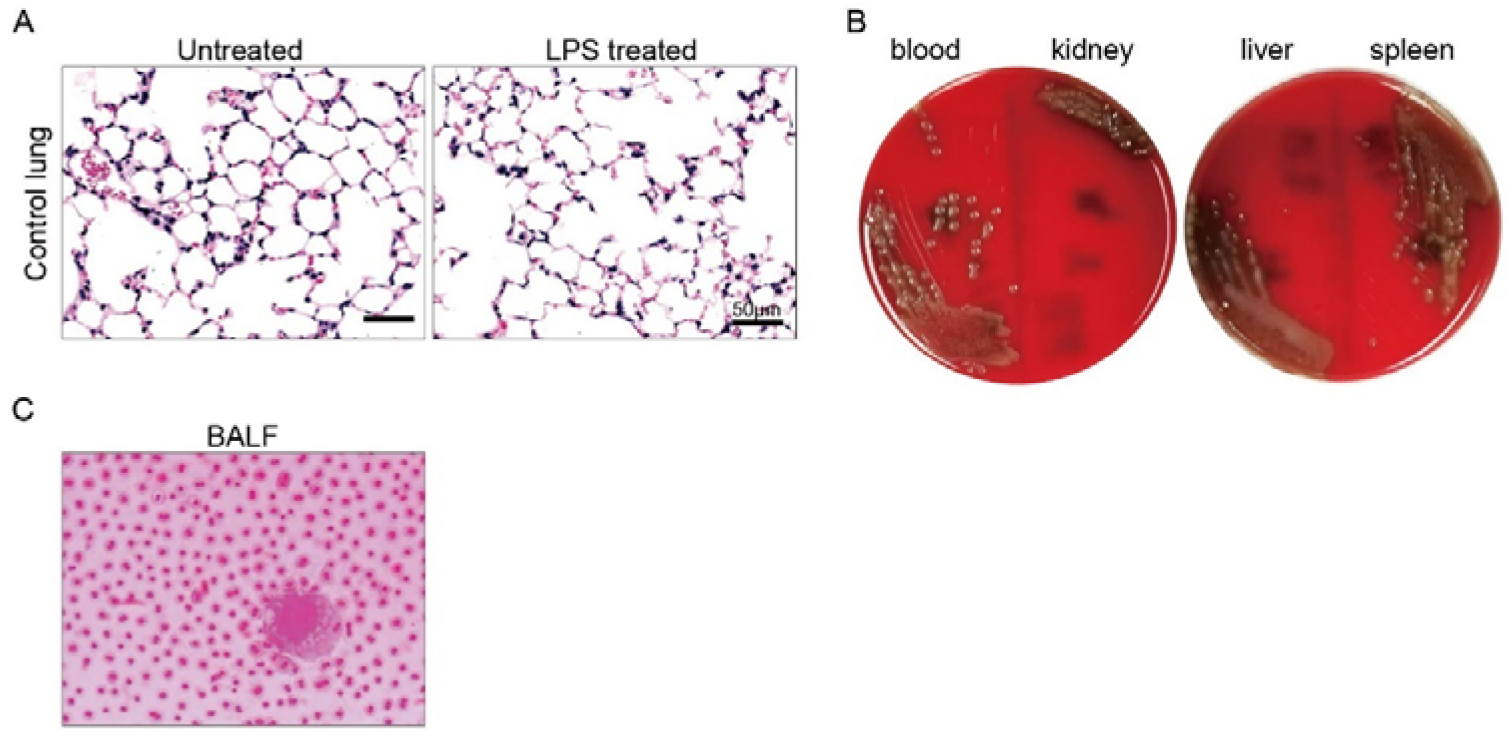
(A) HE stain of lung tissue from the control lung (right lung) from both normal mice (Control) and LPS unilateral instilled mice. No visible difference observed. (B) Tissue culture of mice infected with highly virulent *S. pneumoniae*. All specimen contained high load of *S. pneumoniae*. (C) Bronchoalveolar lavage fluid (BALF) smear and Gram’s stain, from *K. pneumoniae* infected mice. One leukocyte found, surrounded by countless thick capsular *K. pneumoniae*.

**Figure S2.**
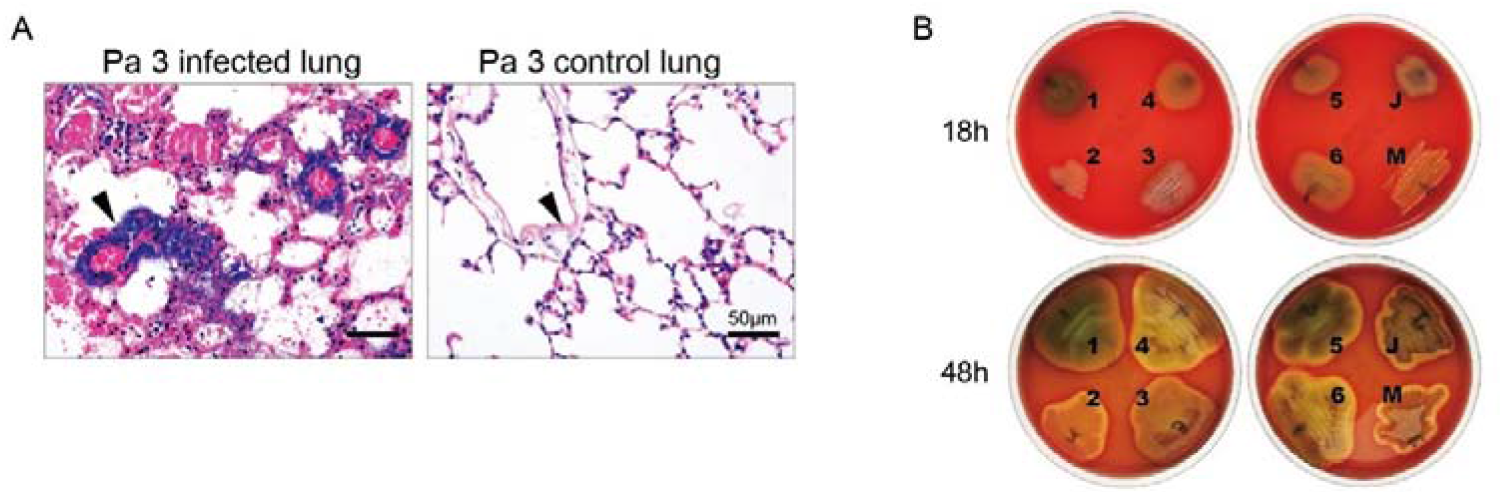
(A) HE stain of strain Pa 3 instilled left lung and un-instilled right lung (control). Bilateral perivascular bacteria proliferation found (black arrowhead). Heavier for the instilled lung. (B) Morphology of 8 *P. aeruginosa* strains cultured on sheep blood agar plates.

**Figure S3.**
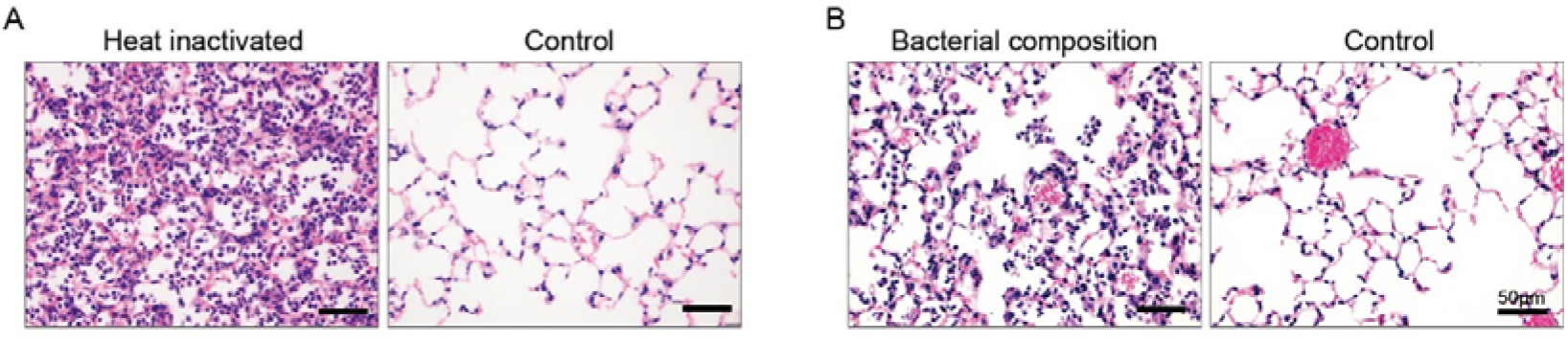
(A) Heated *P. aeruginosa* exoproducts (10 min, 100 ℃) induced neutrophil accumulation, but didn’t induce hemorrhage or alveolar destruction. (B) *P. aeruginosa* bacterial composition induced mild neutrophil infiltration but no other histological change. HE stain.

**Table S1.**
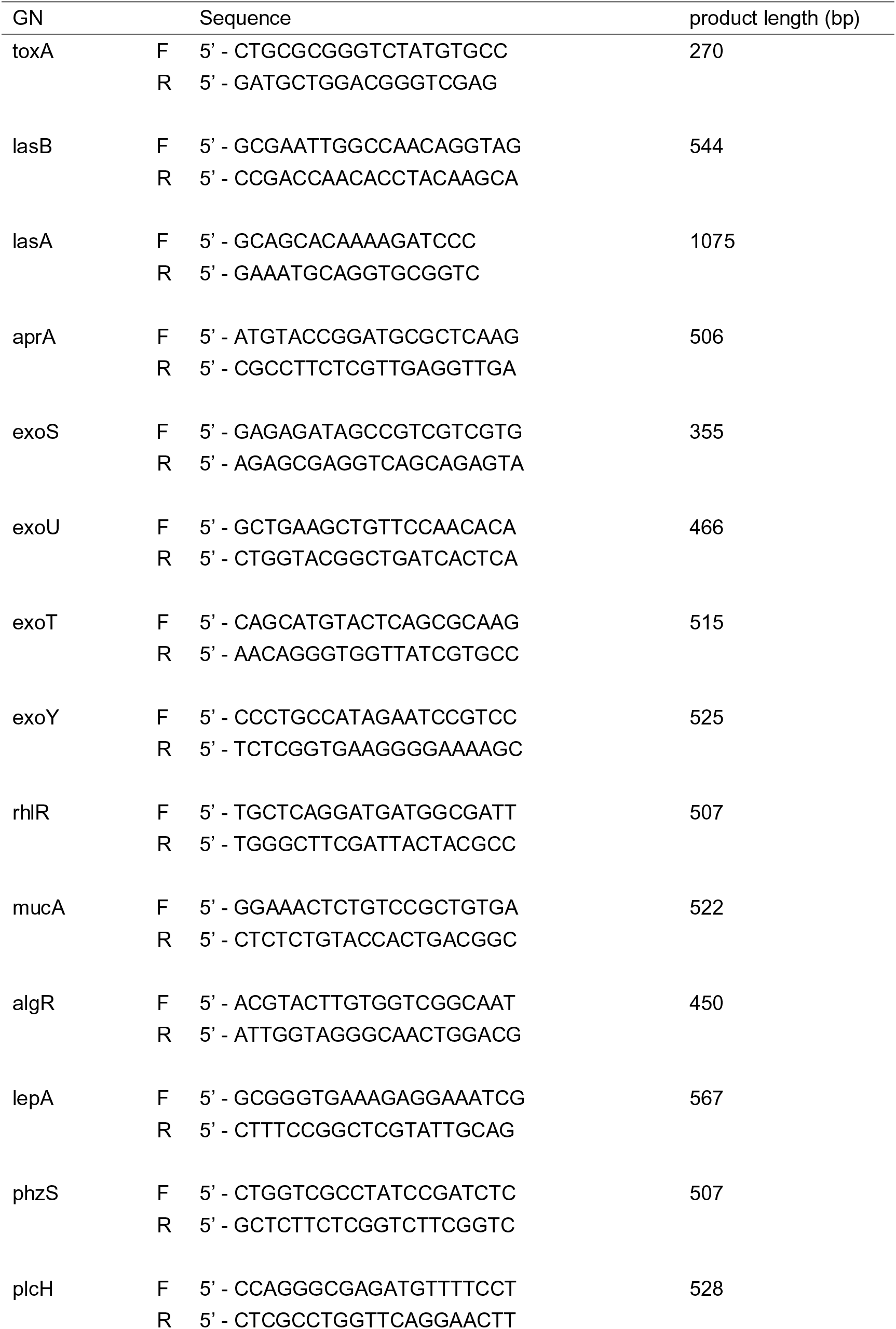
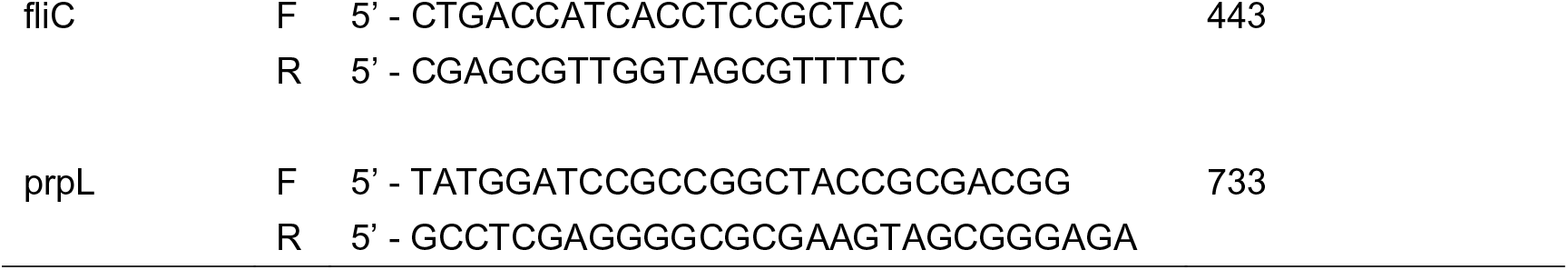
Primer sequences used in PCR for detecting virulence genes.

